# High-frequency stimulation of ventral CA1 neurons reduces amygdala activity and inhibits fear

**DOI:** 10.1101/2020.07.01.183210

**Authors:** Jalina Graham, Alexa D’Ambra, Se Jung Jung, Nina Vishwakarma, Rasika Venkatesh, Abhijna Parigi, Evan G. Antzoulatos, Diasynou Fioravante, Brian J. Wiltgen

**Author notes:** These authors share senior authorship (Diasynou Fioravante, Brian J. Wiltgen). Correspondence should be addressed to: Brian Wiltgen.

## Abstract

The hippocampus can be divided into distinct segments that make unique contributions to learning and memory. The dorsal hippocampus supports cognitive processes like spatial learning and navigation while the ventral hippocampus regulates emotional behaviors related to fear, anxiety and reward. In the current study, we determined how pyramidal cells in ventral CA1 respond to spatial cues and aversive stimulation during a context fear conditioning task. We also examined the effects of high and low frequency stimulation of these neurons on defensive behaviors. Similar to previous work in the dorsal hippocampus, we found that cells in ventral CA1 expressed high-levels of c-Fos in response to a novel spatial environment. Surprisingly, however, the number of activated neurons did not increase when the environment was subsequently paired with footshock. This was true even in the subpopulation of ventral CA1 pyramidal cells that send direct projections to the amygdala. When these cells were stimulated at high-frequencies (20-Hz), we observed feedforward inhibition of basal amygdala neurons and impaired expression of context fear. In contrast, low-frequency stimulation (4-Hz) did not inhibit principal cells in the amygdala and produced a slight increase in fear generalization. Similar results have been reported in dorsal CA1. Therefore, despite the clear differences between the dorsal and ventral hippocampus, CA1 neurons in each segment appear to make similar contributions to context fear conditioning.

## 1 Introduction

The hippocampus can be divided into distinct segments that make unique contributions to learning and memory (Fanselow and Dong 2010). The dorsal hippocampus supports cognitive processes like spatial learning and navigation via interactions with the entorhinal, parahippocampal and retrosplenial cortices (Cenquizca and Swanson 2006, 2007; Strange et al. 2014; Moser and Moser 1998; Moser et al. 2017). The ventral hippocampus (VH), in contrast, regulates emotional behavior through its connections with the amygdala, nucleus accumbens, lateral hypothalamus, BNST and medial prefrontal cortex (Cenquizca and Swanson 2006, 2007; Jimenez et al. 2018; Hoover and Vertes 2007). However, despite these differences, the dorsal and ventral hippocampus share some important properties. They have the same basic architecture (DG, CA3, CA2, CA1) and intrinsic organization (tri-synaptic loop) and neurons in both regions respond to spatial cues (e.g. place cells) (Strange et al. 2014; Kjelstrup et al. 2008; Ishizuka et al. 1990). These parallels suggest that similar computations may be carried out in the DH and VH during cognitive and emotional learning.

Integrating spatial and emotional information depends on interactions between the DH and VH. During context fear conditioning, for example, animals learn to associate a novel environment with aversive footshock. Encoding this relationship requires spatial information from the DH to be transmitted to the basolateral nucleus of the amygdala via the VH (Fanselow and Dong 2010; Xu et al. 2016). However, neurons in the VH do not act as passive relays; their activity is strongly modulated by emotional states like fear and anxiety, which is not the case in the DH (Jimenez et al. 2018; Ciocchi et al. 2015). Consistent with this fact, lesions of the VH reduce stress hormone release and anxiety-related behaviors while damage to the DH does not (Kjelstrup et al. 2002). Place cells in the VH are also distinct; they have large, overlapping place fields that encode behaviorally-relevant contexts as opposed to precise spatial locations (Komorowski et al. 2013). Based on these findings, we hypothesized that dorsal and ventral CA1 neurons would respond to different stimuli during context fear conditioning. Specifically, neurons in dCA1 should respond primarily to the spatial context while cells in vCA1 should be more responsive to footshock.

To examine our hypothesis, we quantified immediate-early gene expression (IEG) in ventral CA1 neurons after spatial exploration and emotional learning. For the former, mice were exposed to a novel environment and for the latter, mice underwent context fear conditioning. We found that ventral CA1 was strongly activated by the new environment but, surprisingly, c-Fos expression did not increase further when the context was paired with shock. Neurons in dorsal CA1 have been shown to respond in the same way under similar conditions (Radulovic et al. 1998; Lovett-Barron et al. 2014). Next, we stimulated ventral CA1 neurons that project to the basal amygdala to determine if defensive behaviors could be induced after context fear conditioning. We found that high-frequency stimulation (20-Hz) disrupted freezing and led to feed-forward inhibition of principal cells in the basal amygdala. In contrast, low frequency stimulation (4-Hz) slightly increased defensive behaviors and did not inhibit the basal amygdala. Similar results have been reported in dorsal CA1 (Ryan et al. 2015). Together, these data suggest that dorsal and ventral CA1 may play similar roles during context fear conditioning despite the functional differences between these subregions.

## Results

### 1.1 Context exploration induces c-Fos expression in vCA1 pyramidal neurons

The dorsal hippocampus responds to spatial and contextual cues while amygdala neurons respond strongly to emotional stimuli like shock (O’Keefe et al. 1971; O’keefe and Speakman 1987; Tanaka et al. 2018; Radulovic et al. 1998; Barot et al. 2009; Beyeler et al. 2018; Pelletier et al. 2005; Wolff et al. 2014). The goal of the current experiment was to determine how ventral CA1(vCA1) neurons respond to these same stimuli. To do this, we compared c-Fos expression after exploration of a novel environment to that observed after context fear conditioning. Expression was quantified in vCA1 pyramidal cells or in the specific neurons that send projections to the basal nucleus of the amygdala (BA). To identify the latter, the retrograde tracer ctb-647 was infused into the BA 3-days prior to training (**Figure 1A, left**). On the training day, Control mice were left in their home cage, the Context group explored a novel environment for 5 minutes and the Context + shock mice underwent contextual fear conditioning (**Figure 1A, middle**). Fear conditioning consisted of two foot-shocks (2 sec, 0.3mA, 1-minute interstimulus-interval) that were delivered after a three-minute exploration period. Ninety-minutes after training, the animals were sacrificed, and their brains fixed for c-Fos immunohistochemistry.

**Figure 1.**
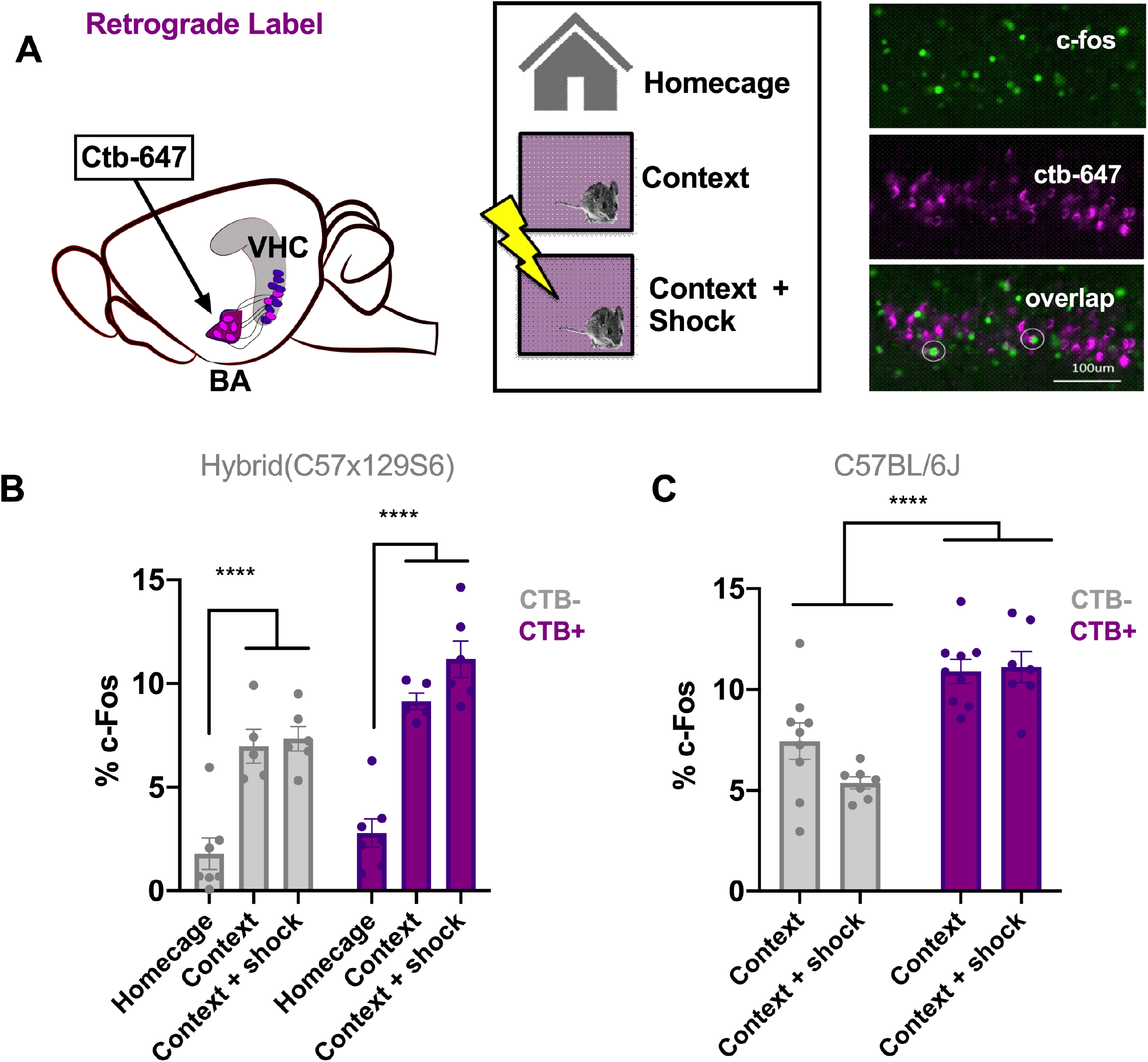
Context exploration induces c-Fos expression in vCA1 pyramidal neurons. **A**. Experiment design: Ctb was injected into the basal amygdala **(left)**. Animals were divided into three groups after surgery/handling **(middle)**. The ‘context’ group was allowed to explore the context for six minutes and the ‘context+shock’ group received 3 mild shocks (0.6mA, 1 min ISI) in the context after 3 minutes of exploration. The control group was left in the homecage. Animals were sacrificed 90 minutes after the start of their behavior session and the brain tissue was fixed for c-Fos immunostaining. **(right)** shows an example of c-Fos staining in green, Ctb labeling in magenta and overlap indicating activation of projection specific neurons. **B**. The percent of Ctb+ and Ctb-neurons which respond to control, context or context + shock conditions. Exploration of the context alone or experiencing context fear learning resulted in increased c-Fos expression relative to controls. There was no difference between context and context + shock groups. **C**. The percent of ctb+ and ctb-neurons in BA-projecting vCA1 neurons of C57 mice. There was no difference between context and context + shock conditions. All data are presented as mean ± SEM and * indicates a significant difference between conditions.

Relative to the Control group, we found a significant increase in the percentage of c-Fos positive neurons after exploration of a novel environment and following context fear conditioning. These increases were similar for the Context and Context + Shock groups and they were observed both in ventral CA1 neurons that project to BA (Ctb+ neurons) as well as those that do not (Ctb-neurons). Interestingly, in all groups, c-Fos was expressed in a larger percentage of Ctb+ cells than Ctb-cells (**Figure 1B**) [Repeated Measures ANOVA, Main effect of Group, F (2, 15) = 42.99, p < .0001; Main effect of Cell type, F (1, 15) = 22.93, p = .0002; No Group x Cell type interaction F (2, 15) = 3.115, p = .0739; Bonferroni post-hoc tests, Control vs Context (p < .0001), Control vs Context + Shock (p < .0001), Context vs Context + shock (p = .5778)].

While c-Fos expression did not increase in Ctb+ vCA1 neurons after context fear conditioning compared to exploration alone, there was a trend in this direction. One difference between our study and previous work is that we used F1 hybrids (129S x C57) and trained them with relatively weak shocks because these mice learn much better than inbred C57s (Balogh and Wehner 2003; Owen et al. 1997). To address this discrepancy, we quantified c-Fos expression in C57Bl/6J mice that either explored the environment or underwent fear conditioning with three, 0.75mA shocks. Similar to the results from our F1 hybrids, CTB+ neurons expressed more c-Fos than CTB-neurons and there were no differences between the exploration and Context + Shock groups [2-way ANOVA, Main effect of cell type(Ctb+/Ctb-) F(1, 28) = 41.37, p < 0.0001; No effect of learning condition F(1, 28) = 1.685, p = 0.2049; No condition x cell type interaction F(1, 28) = 2.552, p = 0.1213].

Together, these experiments suggest that immediate early gene expression in ventral CA1 neurons is strongly induced by exposure to a novel environment as is the case in dorsal CA1 (Radulovic et al. 1998; Barot et al. 2009). The addition of shock did not increase expression relative to context exploration alone as observed in subcortical structures like the amygdala (Radulovic et al. 1998; Barot et al. 2009; Milanovic et al. 1998). It remains possible, however, that context exploration and footshock activate many of the same neurons in vCA1, making it difficult to find a significant increase in the total number of c-Fos+ neurons. This possibility could be addressed in future studies by using tools like catFISH to identify vCA1 neurons that respond to both stimuli (Barot et al. 2009). Fear conditioning may also have changed the excitability and/or firing pattern of vCA1 neurons in a way that did not alter immediate early gene expression.

### 1.2 High frequency stimulation of vCA1 neurons disrupts the expression of context fear

Previous work found that high-frequency stimulation of dCA1 neurons (20-Hz) after fear conditioning did not induce freezing like it does in the dorsal dentate gyrus (DG) (Ryan et al. 2015; Ramirez et al. 2013). This may have been the case because dCA1 does not send a direct projection to the ventral hippocampus like dDG and dCA3 (Fricke and Cowan 1978; Ishizuka et al. 1990; Swanson et al. 1978). To examine this possibility, we performed high-frequency stimulation of vCA1 neurons and determined the effects on freezing behavior in previously fear conditioned mice. To identify optimal stimulation frequencies, we first injected AAV5-ChETA-EYFP into the ventral hippocampi of 3-4 week old mice and extracted the brains for slice electrophysiology 2-3 weeks later. We recorded from ChETA-EYFP-expressing vCA1 neurons using a cell-attached patch configuration (see methods for details) while stimulating with 488 nm light at 10, 20 or 50 Hz (**Figure 2)**. As observed in the dorsal hippocampus, pyramidal cells in vCA1 could easily follow 10 Hz and 20 Hz optogenetic stimulation across multiple trials (**Figure 2D)**. At these frequencies, the spike probability for light pulses 2-5 in a train was close to 1 and not significantly different from the spike probability for light pulse 1 [permutation test for pulses 2-5 compared to pulse 1: **10 Hz**: p = 0.82, p = 0.82, p = 0.82, p = 0.20; **20 Hz**: p = 0.89, p = 0.88, p = 0.62, p = 0.43]. In contrast, when the same neurons were stimulated at 50 Hz, they only responded reliably to the first light pulse (average spike prob. ± SEM: 1.0 ± 0.00). The firing probability to subsequent stimuli progressively decreased, and was significantly reduced by pulse 4(**Figure 2D)**. [permutation test for pulses 2-5 compared to pulse 1: **50 Hz**: p = 0.17, p = 0.09, p = 0.01, p = 0.001]

**Figure 2.**
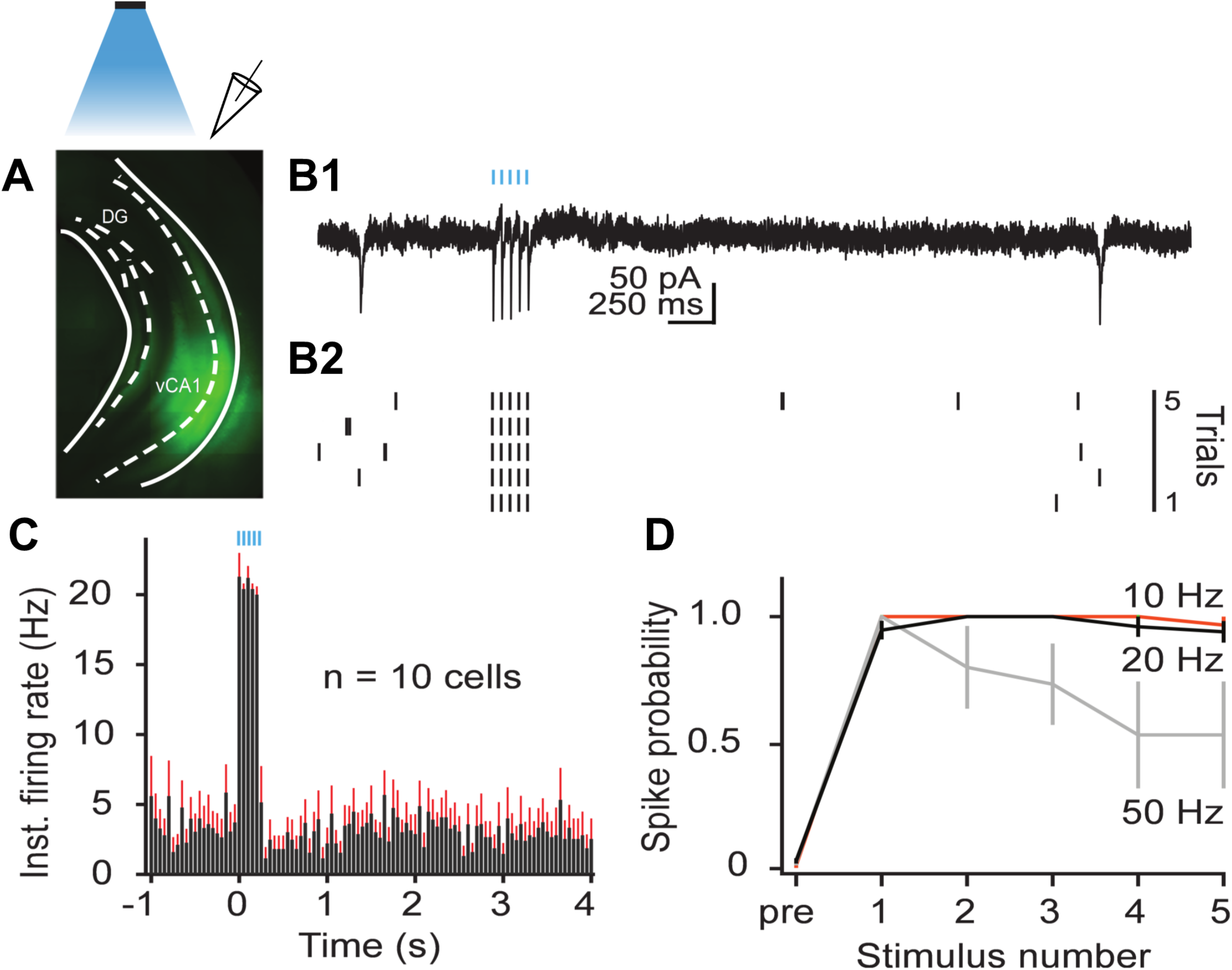
Stimulation of ChETA at 20Hz reliably activates vCA1 pyramidal neurons. **A**. Expression of ChETA-EYFP in ventral/intermediate CA1. **B**. Ex vivo cell-attached recording from a representative ChETA-expressing vCA1 pyramidal neuron in response to 5 10-ms light pulses at 20 Hz (blue lines). **B1**. Single example trial; **B2**. Raster plot of 5 trials. **C**. Average instantaneous firing rate (in spikes/s) for 10 vCA1 pyramindal neurons (bins: 50 ms, black: average firing rate, red: SEM). F, Average spike probability for 3 photostimulation frequencies (10, 20 and 50 Hz).

Based on these results, we decided to stimulate vCA1 pyramidal neurons at 20 Hz after context fear conditioning. Our initial plan was to selectively stimulate c-Fos+ neurons by infusing AAV-TRE-ChETA into the ventral hippocampus of TetTag mice purchased from Jackson Labs. However, we observed a significant amount of non-specific labeling in these mice compared to the original fos-tTA mouse line (Tayler et al. 2011; Tanaka et al. 2014; Nakazawa et al. 2016; Crestani et al 2019; Wilmot et al. 2018). Consequently, instead of tagging memory cells, we selectively stimulated vCA1 neurons that projected to the basal amygdala. To do this, AAVrg-EBFP-Cre was injected into the basal amygdala and FLEX-ChETA-mCherry virus was infused into the ventral hippocampus. Bilateral optic fibers were implanted directly over ventral CA1 (**Figure 3A**). Histological analyses confirmed that ChETA-mCherry expression was expressed in ventral CA1 (**Figure 3B, top**) and the highest density of Cre expression associated with the injection site was restricted to basal amygdala (**Figure 3B, bottom**). A second group received infusions of CaMKII-ChETA-EYFP into ventral CA1 to examine the effects of non-selective stimulation on freezing. Control groups received infusions of AAV-CaMKII-EYFP (n=4) into ventralCA1 or combined injections of AAVrg-EBFP-Cre into basal amygdala and FLEX-tdTomato in ventral CA1 (n=4).

**Figure 3.**
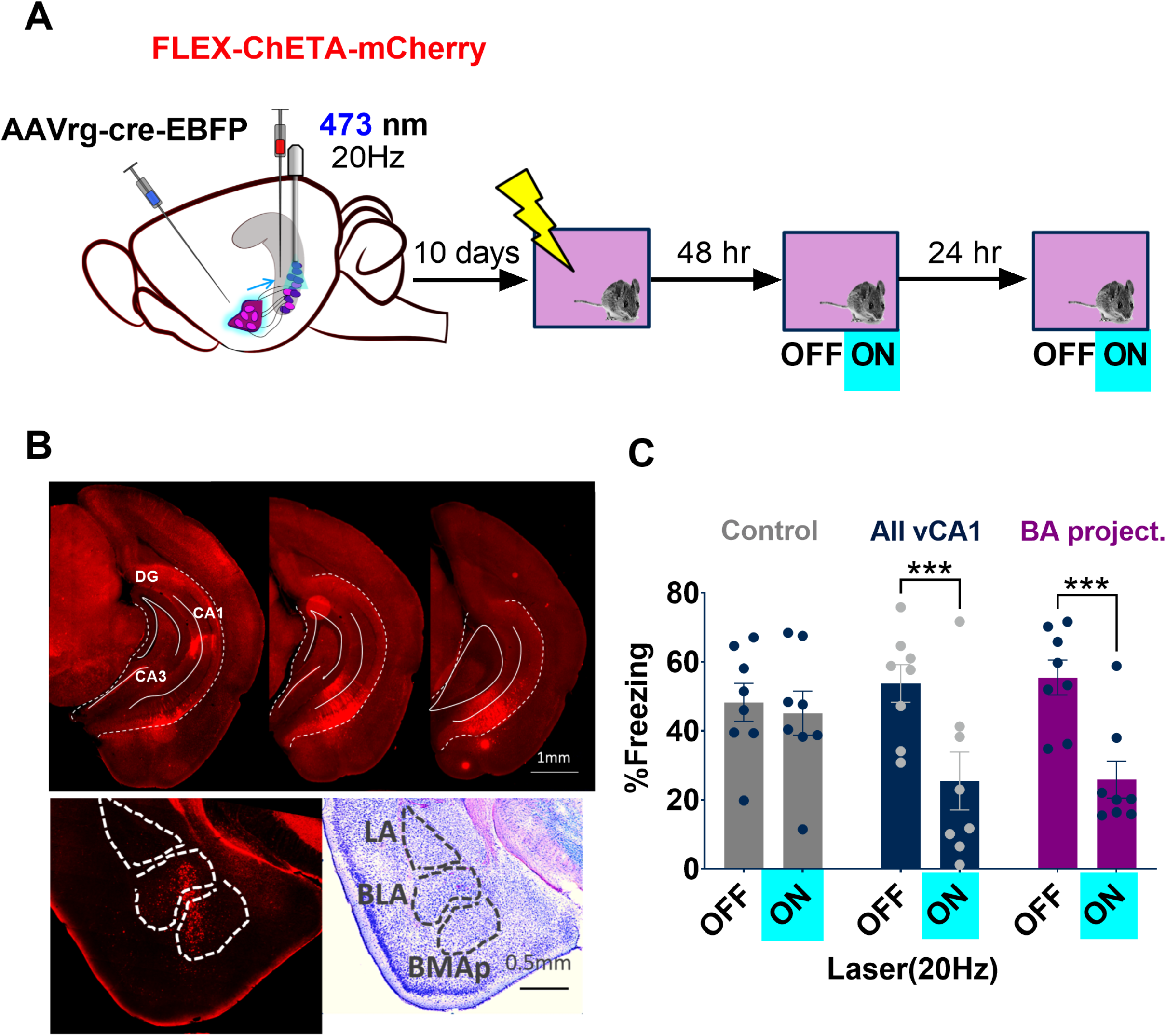
High frequency stimulation of vCA1 neurons disrupts the expression of context fear. **A**. Experiment design. ChETA expression was driven in the VHC-BA projection (left) using a combination of a retrograde cre virus in the BA (AAVrg-pmsyn1-EBFP-cre) and a cre dependent virus in ventral hippocampus(AAV5-EF1a-DIO-ChETA-mCherry). An optic fiber was implanted above ventral CA1. For nonspecific ‘All vCA1’-stimulation groups, CaMKII-ChETA-EYFP was injected into the VHC. One half of the controls underwent the same surgery as the projection specific group, with the substitution of the FLEX ChETA virus for a FLEX-EYFP and one half had CaMKII-EYFP injected into VHC. After recovery, animals were trained as described in the methods and placed into the training context for memory retrieval test 48 and 72 hours later. Each test session started with three minutes baseline (BL) and ended with three minutes of laser stimulation (473 nm, 20Hz, 15 msec pulse duration). **B**. (top) Expression of FLEX-ChETA-mCherry in ventral CA1. (bottom left), Immunostaining for cre reveals the injection site of AAVrg-EBFP-cre. Injection site location was confirmed by identifying amygdala nuclei on adjacent nissl sections (bottom right). **C**. Average freezing during baseline and stimulation epochs of both memory retrieval test days. During the baseline, all groups froze at similar levels, indicating successful acquisition of fear memories by both controls and ChETA expressing animals. However, when vCA1 was stimulated, freezing was reduced in both whole VHC and VHC-BA projecting ChETA groups relative to controls. All data are presented as mean ± SEM and * indicates a significant difference between conditions.

Following recovery from surgery, animals were handled and habituated to the optic fiber cable for 5 days and then underwent context fear conditioning. Training consisted of a 3-min baseline period followed by 2 shocks (0.3 mA, 2 sec duration) delivered 1 minute apart. Two days later, mice were placed back in the training environment to assess context fear memory. The test began with a 3-minute baseline period followed by 3 minutes of blue light stimulation (473, 10 mW, 20 Hz). Mice received an identical test 24 hours later (**Figure 3A**).

During the baseline period, freezing was similar for all groups (**Figure 3C**). However, during 20-Hz laser stimulation, both ChETA-expressing groups exhibited a significant decrease in freezing that was not observed in EYFP controls. [Repeated measures 2-way ANOVA, Stimulation x Group interaction (F (2, 21) = 5.88, p = .0093; Bonferroni post-hoc tests multiplicity adjusted p-values, Control On vs Off (p >0.9999), All vCA1 On vs Off (p = 0.0005) BA- projecting vCA1 On vs Off (p =0.0003)]. The size of this deficit was similar whether all excitatory ventral CA1 neurons were stimulated or just those that project to the basal amygdala. No Group x laser interaction was observed [2-way ANOVA F(1,14) = 0.01590, p = 0.9014]. In a separate group of animals, we confirmed that laser stimulation activated ventral CA1 pyramidal neurons and did not likely affect basal amygdala cells directly given the location of our fibers and the light intensity used to activate ChETA (**Supplementary figure 1**). Therefore, high-frequency stimulation of vCA1 neurons after fear conditioning does not increase freezing, similar to results obtained in dCA1(Ramirez et al.2013; Wilmot et al. 2019; Krueger et al. in press).

### 1.3 High frequency stimulation of vCA1 terminals inhibits principal cells in the BA

Stimulation of vCA1 neurons at high-frequencies can produce feed-forward inhibition of principal cells in the basal amygdala (Hübner et al. 2014; Bazelot et al. 2015). To examine this possibility, we recorded from basal amygdala neurons while stimulating ventral CA1 terminals at high (20 Hz) or low (4 Hz) frequencies. AAV5-CaMKII-ChETA was infused into the ventral hippocampus and coronal slices were taken from the BA 2-3weeks later. BA neurons were excited by 4 Hz stimulation of vCA1 terminals and fired action potentials to every light pulse in a stimulus train (average spike prob ± SEM: pulse 1: 0.97 ± 0.02; pulse 2: 0.88 ± 0.06, pulse 3: 0.86 ± 0.05, pulse 4: 0.86 ± 0.05, pulse 5: 0.91 ± 0.04). In contrast, 20 Hz stimulation produced only a single action potential and suppressed responding to all remaining light pulses in a train of 5 pulses (average spike prob. ± SEM: pulse 1: 0.95 ± 0.02; pulse 2: 0.06 ± 0.03, pulse 3: 0.01 ± 0.01, pulse 4: 0.05 ± 0.04, pulse 5: 0.04 ± 0.03) [permutation test, pulses 1-5, **4 Hz** vs **20 Hz**: p = 0.09, p = 0.00, p = 0.00, p = 0.00, p = 0.00] (**Figure 4C**). To determine if this suppression was caused by local inhibition, we stimulated vCA1 terminals at 20 Hz in the presence of the GABA_A_ receptor antagonist bicuculine. This manipulation partially rescued the activity of BA neurons (average spike prob. ± SEM: pulse 1: 0.96 ± 0.04; pulse 2: 0.72 ± 0.08, pulse 3: 0.25 ± 0.11, pulse 4: 0.09 ± 0.09, pulse 5: 0.05 ± 0.05) [permutation test, pulses 1-5, **20 Hz** vs **20 Hz** + **Bic**: p = 0.59, p = 0.001, p = 0.18, p = 0.38, p = 0.48], suggesting that feed-forward inhibition plays a role in suppressing excitatory activity when vCA1 neurons are stimulated at high frequencies (Bazelot et al. 2015; Hübner et al. 2014). Given that firing was not completely rescued, other factors like synaptic depression are likely to contribute to this effect.

**Figure 4.**
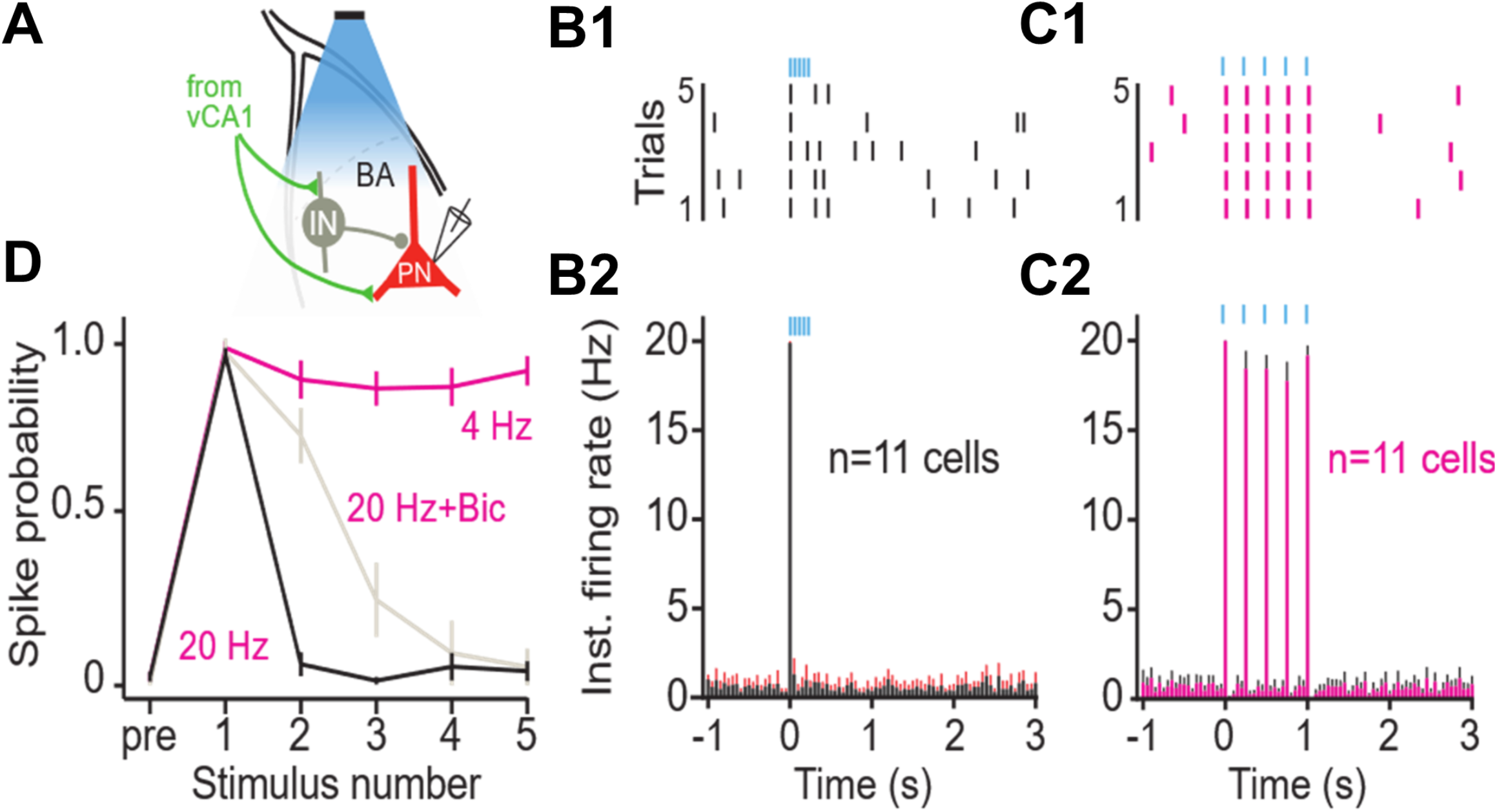
High frequency stimulation of vCA1 terminals inhibits principal cells in the BA. **A**. Schematic of the experimental configuration: vCA1 axons expressing ChETA-EYFP were optically stimulated in the BA while recording on-cell from BA PNs. IN: interneuron. **B-C**. Example raster plots (**B1-C1**) and average instantaneous firing rates (+SEM; **B2-C2**) for BA PNs in response to 5 10-ms light pulses at 20 (**B**) and 4 (**C**) Hz (bins: 50 ms). **D**. Average spike probability for 4 Hz, 20 Hz, and 20 Hz in the presence of the GABAAR antagonist bicuculine (Bic).

### 1.4 Low frequency stimulation of vCA1 does not disrupt the expression of context fear

Activity in the hippocampus, amygdala and prefrontal cortex is synchronized around theta-frequency (4-12 Hz) during aversive learning, consolidation and fear expression (Seidenbecher et al. 2003; Padilla-Coreano et al. 2016; Narayanan et al. 2007; Lesting et al. 2011). Given that BA neurons can follow when vCA1 terminals are stimulated at 4 Hz, we examined the impact of this manipulation on the expression of fear. Mice received bilateral infusions of AAV-CaMKII-ChETA-EYFP (n = 6) or AAV-CaMKII-EYFP (n = 5) into the ventral hippocampus and optic fibers were implanted above vCA1. Two weeks later, they were trained on context fear conditioning as described above. Memory was tested 48 hours after training and vCA1 neurons were stimulated at 4 Hz (10 mW, 4 Hz, 15 msec pulses) during the last 3 minutes of the session (**Figure 5A**). Unlike high-frequency stimulation, this manipulation did not disrupt the expression of context fear (**Figure 5B**) [2-way ANOVA, no laser x virus interaction F = 0.05282, p = 0.824, no main effect of laser F(1,9) = 0.005282, p = 0.8234].

**Figure 5.**
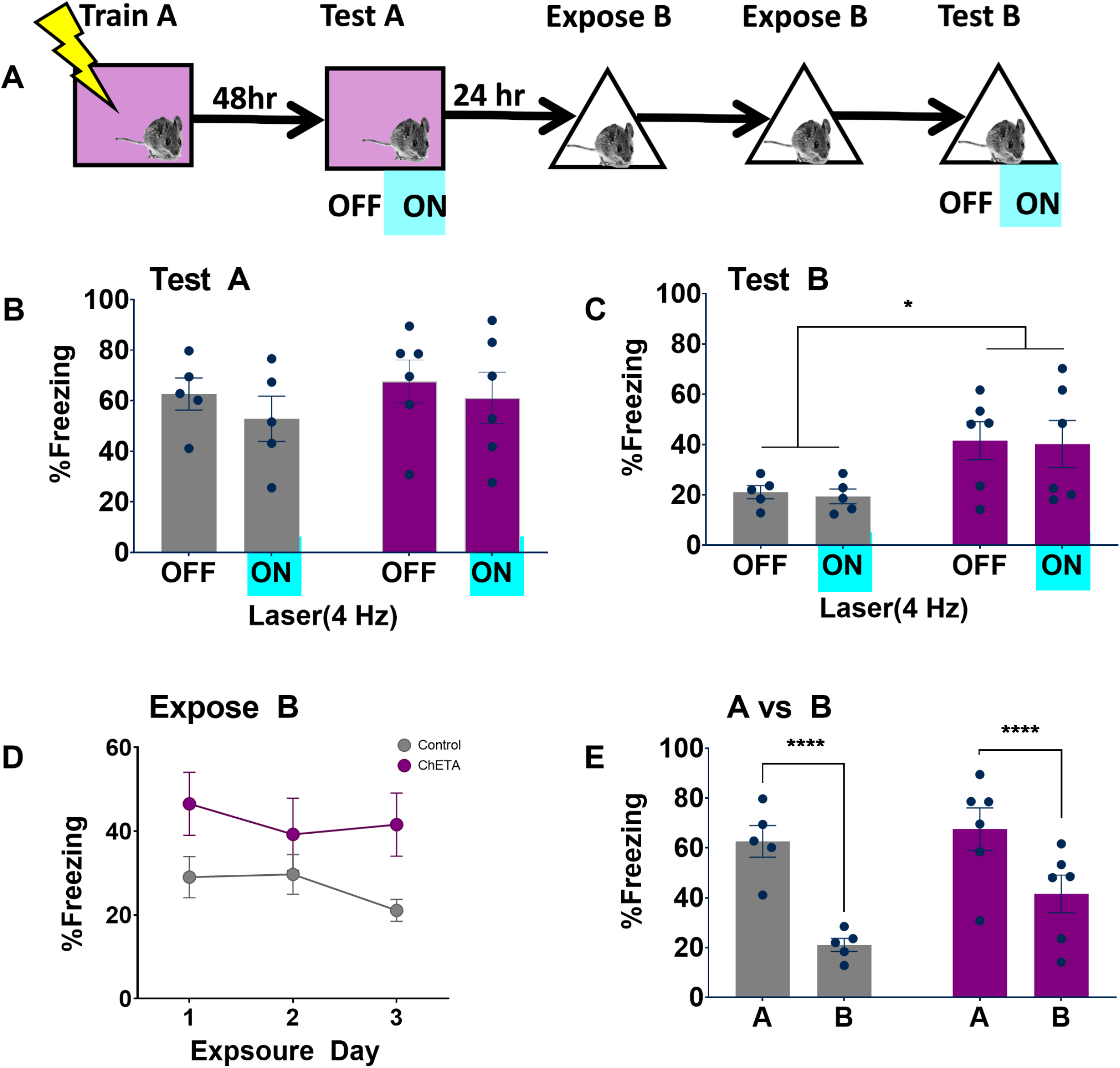
Low frequency stimulation of vCA1 neurons does not disrupt the expression of context fear. **A**. 4Hz stimulation during the last three minutes of a six-minute fear memory test in the training environment (context A) did not affect freezing in the ChETA group (blue) relative to the EYFP controls (grey). **B. Mice stimulated at** 4Hz stimulation in context A, exhibited more fear generalization to a novel environment (context B) than control animals. However, stimulation in context B did not lead to additional increases in fear. **C**. 4Hz stimulation in context A did not alter extinction of generalized fear to context (B). **D**. 4Hz stimulation did not significantly alter discrimination between context A and context B. All data are presented as mean ± SEM and * indicates a significant difference between conditions.

### 1.5 Low frequency stimulation of vCA1 neurons does not induce freezing but increases fear generalization

We next determined if 4Hz stimulation could induce freezing in a novel environment (context B). The same mice were pre-exposed to context B for 2 days to extinguish any generalized fear **(Figure 5A)**. On day 3, the animals were put back into context B and vCA1 neurons were stimulated at 4 Hz during the last 3-minutes of the session. This manipulation did not increase freezing in ChETA mice relative to control animals [2-way repeated measures ANOVA, no Group x Laser interaction, F (1,9) = .001, p = 0.968]. However, previously stimulated ChETA mice did exhibit an overall increase in freezing during the test session [Main effect of Group, F (1,9) = 5.50, p = .043]. This suggests that vCA1 stimulation in context A may have increased generalization to context B or reduced fear extinction during the pre-exposure sessions. To examine these possibilities, we analyzed freezing during the first 3-minutes of each session in context B (i.e. laser off periods) (**Figure 5D)**. During these sessions, there was a trend for increased freezing in ChETA mice, but it did not reach statistical significance. Context freezing did not decrease significantly across pre-sessions in either group, indicating that they were not sufficient to produce robust extinction [2-way repeated measures ANOVA, No Group x Session interaction F (2,18) = 1.38, p = 0.27; No effect of Group F (1,9) = 3.31, p = 0.1; No effect of session, F (2, 18) = 1.81, p = .19].

To examine discrimination, we compared freezing during the first 3-minutes of the test session in context A to the first 3-minutes of the initial context B exposure session (**Figure 5E)**. Both ChETA and control groups showed significantly more freezing to the training context [Two way ANOVA main effect of context F (1, 9) = 76.30, p < 0.0001]. There was a trend for reduced discrimination in ChETA mice, but it failed to reach statistical significance [2-way repeated measures ANOVA, No group x context interaction F (1,9) = 4.04, p = .075; no effect of group F (1,9) = 1.94, p = .19]. We should note, however, that our behavioral procedures were designed to observe increases in freezing when vCA1 neurons were stimulated, not to examine changes in context discrimination or extinction. A more thorough analysis of these processes will be required to determine if they are affected by low-frequency stimulation of vCA1 neurons.

## 2 Discussion

During context fear conditioning, spatial information is transmitted from dorsal to ventral hippocampus and then relayed to the amygdala where it is associated with shock (Wiltgen et al. 2006; Sutherland et al. 2008; Maren and Fanselow 1995; Xu et al. 2016; Jimenez et al. 2018; Kim and Cho 2020). We examined the relationship between fear expression and activity in the ventral hippocampus to amygdala pathway. Similar to results obtained in the dorsal hippocampus, we found that c-Fos expression increased significantly in vCA1 neurons after exposure to a novel environment (Radulovic et al. 1998). However, the addition of shock did not alter the number of labelled cells as it does in other structures like the amygdala (Barot et al. 2009; Milanovic et al., 1998; Radulovic et al. 1998). This result was somewhat surprising given the role of the ventral hippocampus in fear and anxiety and its strong connectivity with subcortical regions that are involved in emotion (Jimenez et al. 2018; Hoover and Vertes 2007; Cenquizca and Swanson 2006, 2007). However, a recent study found that vCA1 neurons activated during exploration are the same cells that increase their connectivity with basal amygdala neurons after the environment is paired with shock (Kim and Cho 2020). Together with the current results, these data suggest that the ventral hippocampus may primarily transmit context information to the amygdala during fear conditioning.

High-frequency stimulation (20 Hz) of “engram cells” in dorsal DG and CA3 has been shown to induce freezing after context fear conditioning (Ryan et al. 2015; Ramirez et al. 2013). However, the same manipulation does not drive freezing in dorsal CA1 (Ryan et al. 2015). This may be the case because dCA1 does not send a direct projection to the ventral hippocampus like dDG and dCA3 (Fricke and Cowan 1978; Swanson et al. 1978; Ishizuka et al. 1990). However, when we stimulated vCA1 neurons that project to the BA at 20 Hz, freezing was significantly impaired. This finding is similar to a recent report that showed 10-Hz stimulation of vCA1 terminals in the BA impairs the expression of context fear (Jimenez et al. 2018). To determine why high-frequency stimulation was disruptive, we recorded from principal cells in the BA while activating vCA1 terminals. We found that high-frequency stimulation (20 Hz) inhibited excitatory activity in BA neurons while low-frequency stimulation (4Hz) did not. The inhibitory effect was reduced by a GABA antagonist, indicating that it is due, in part, to feed-forward inhibition. Interestingly, the medial prefrontal cortex (mPFC) sends a long-range projection to the BA that could disinhibit principal cells and allow them to respond to strong input from vCA1 (Hübner et al. 2014). A circuit like this would be ideal for selecting adaptive responses in different situations. For example, place cell activity in vCA1 could induce exploration or freezing in the same environment depending on the input from mPFC. Future work can examine this idea *in vivo* by manipulating both the ventral hippocampal and mPFC inputs to the BA.

Unlike high-frequency stimulation, 4 Hz activation of vCA1 neurons did not inhibit principal cells in the BA. Given that the hippocampus, amygdala and mPFC all oscillate at theta-frequency during fear expression, we hypothesized that low-frequency stimulation may enhance freezing rather than impair it (Seidenbecher et al. 2003; Padilla-Coreano et al. 2016; Narayanan et al. 2007; Lesting et al. 2011; Courtin et al. 2014; Karalis et al. 2016). Consistent with this idea, 4 Hz stimulation did not disrupt freezing and produced a slight increase in fear generalization. However, freezing did not increase during laser stimulation itself, which has been observed when “engram cells” in dCA1 are stimulated at 4 Hz (Ryan et al. 2015). Therefore, our behavioral effects may be small because we did not selectively stimulate vCA1 neurons that were activated during training. These “engram cells” were recently shown to increase their connectivity with BA neurons after context fear conditioning (Kim and Cho 2020). It is possible, therefore, that low-frequency stimulation of vCA1 “engram cells” would enhance freezing after training. High-frequency stimulation may also be able to induce fear when it co-occurs with mPFC-mediated disinhibition of the BA. This pathway has been shown to disinhibit principal cells in the BA and drive freezing, but it is not known if it mediates context fear by gating excitatory input from vCA1 (Hübner et al. 2014; Karalis et al. 2016). Additional research is needed to examine these possibilities.

To summarize, our results indicate that neurons in dorsal and ventral CA1 share some important similarities despite the distinct connectivity and function of these subregions. Cells in each area express high levels of c-Fos in response to a novel context and the size of this effect does not change if the environment is paired with shock. High-frequency stimulation of pyramidal cells in either subregion also disrupts the expression of freezing after context fear conditioning. Low-frequency stimulation, in contrast, induces freezing in dCA1 engram cells and slightly increases fear generalization in vCA1. Additional research will be required to identify the specific patterns of activity that drive freezing in vCA1 and determine how they interact with input from other brain regions like the mPFC.

## Methods

### 2.1 Subjects

Experiments in Figures 1, 3, and 5 used hybrid mice generated at Taconic from C57BL/6NTac crossing female mice with 129S6/SvEvTac male mice from Taconic’s US’s commercial colonies **(IMSR Cat# TAC:b6129, RRID:IMSR_TAC:b6129)**. **Figure 1C** used male C57Bl/6J mice from the Jackson Laboratory (**IMSR Cat# JAX:000664, RRID:IMSR_JAX:000664**). For ease of use, the electrophysiology experiments (Figures 2 and 4) and the activation experiment shown in Supplementary Figure 1 used male and female wildtype C57BL/6J mice from our breeding colony. Animals were maintained on a 12 hour light-dark cycle and all surgical and behavioral procedures took place during the light phase. All animals had ad libitum access to food and water. All animals were single housed post-surgery. All experiments were approved by UC Davis Institutional Animal Care and Use (IACUC).

### 2.2 Surgeries

Mice 9-20 weeks old (age range smaller for individual experiments) were induced at 5% and maintained on 1-3% isoflurane mixed with O2. A feedback-controlled heating pad was used to maintain body temperature as 37°C throughout surgery. Small holes were drilled above the injection sites targeting either bilateral VHC(AP, −3 mm; ML, ±3.5 mm; DV −3.9 & −3.5 mm from dura, 250nl each site) BA(AP, −1.55 mm; ML, ± 2.85; DV, −5/-4.8 mm from dura), or both. For the electrophysiology experiments, mice were 3-4 weeks old at the time of surgery and coordinates and volume were adjusted for size (AP, −2.8 mm; ML, ±3.6 mm; DV, −2.8 mm from brain surface, 350 nl virus). Note: for the AAVrg experiments, the most dorsal AMY coordinate was changed to −4.9 mm from dura due to increased spread of the virus (37nl/site). Ctb experiments used 50 nl of Ctb-647(Invitrogen) per site. For optogenetic experiments, an additional 3 small holes were drilled for flat-bottomed, stainless steel jewelers’ screws to allow the implantation of optic fiber cannulas. A new mineral oil filled glass pipette with tip diameter between 25-40 μm was used to withdraw and inject virus or tracer per animal. All virus or tracer was injected per DV site at 2 nl sec^−1^ bilaterally at each site. We waited for 3-5 minutes after injection at each site before withdrawing the pipette. Virus volume and rate was precisely controlled using a World Precision Instruments pump (model UMP3). Optic fiber cannulas(0.39 NA, 200 μm diameter, Thorlabs) were manufactured as previously described(Sparta 2012) and implanted bilaterally above virus injection site in ventral CA1 (AP, −3 mm; ML, ±3.75 mm, length 3.4 mm) using Bosworth Trim II dental acrylic to secure the implants to the skull.

### 2.3 Contextual fear conditioning and optogenetic stimulation

Stereotaxic surgery was performed on day 1. On Day 8-12, all animals were handled for 5 minutes a day (either in the vivarium or in a room adjacent to the fear conditioning chambers). Animals in optogenetic experiments had their implants attached to a 1-m split optic patch cable (0.22 NA, 200μm diameter) during handling. All mice excepting those in tagging experiments were trained 24 hours after the last handling session. Training consisted of 3 minutes of context exploration and either 2 shocks (0.3 mA, 1 min ITI, Taconic hybrids) or 3 shocks (0.6mA, 1 min ITI, C57s). Testing consisted of 5-10 minutes in the training context or a novel context (context B). Mice were sacrificed after their final testing session or training (ctb experiment) and c-fos was quantified as described below. For optogenetic experiments, a 473 nm, 300MW DPSS laser system (OptoEngine) was coupled to the branched optic cable and implant through a rotating comutator fixed above the conditioning chamber. Laser output was adjusted to obtain 10 mW from the optic fiber tip measured with an optical power meter (Thorlabs) before each experiment. Doric’s OptG4 software was used to control laser pulse frequency and Med Associates SG-231 28V DC-TTL adapter was used to control onset and offset of laser pulses during behavioral sessions. Laser stimulation consisted of 3 minute epochs with 15 msec pulses at 20 Hz. Mice were trained and tested in Med Associates fear conditioning chambers (30.5 cm x 24.1 cm x 21.0 cm) that were housed in sound-attenuating boxes. Each chamber consisted of a stainless steel grid floor, overhead LED lighting and a scanning, charge-coupled device video camera. Context A consisted of the grid floor, no wall inserts and cleaned with 70% EtOH. Context B consisted of a smooth plastic floor insert, a curved plastic wall insert, 1.5 pinches of clean corncob bedding and cleaned with PDI Sani Cloth Plus germicidal wipes.

### 2.4 Immunohistochemistry and microscopic imaging

Animals were sacrificed 90 minutes after training, testing, or final laser stimulation. Mice were deeply anesthetized using 5% isoflurane mixed with O2 and then transcardially perfused first with 0.1M Phosphate buffered saline(PBS) and fixed with 4% paraformaldehyde(PFA). Brains were extracted and left in PFA for 24-48 hours and then sliced into 40 μm coronal sections using a Leica VT-1000 vibratome. To visualize virus spread and locate injection sites and fiber optic tips, two separate series spanning the anterior-posterior axis (every 5^th^ and 6^th^ slice). For Ctb injection site localization, one series was Nissl-stained and amygdala nuclei were identified using the online Allen interactive mouse reference atlas. The adjacent sections in the other series were stained with DAPI and the location of the injection site was mapped onto the identified Amygdala nuclei. Slices 1-4 were stored for later c-fos immunohistochemistry. 3-4 sections per area were randomly chosen for cfos quantification. Slices were incubated in blocking buffer (2% normal Donkey serum, 0.2% Triton-X 100 in 0.1 M PBS) for 15 minutes followed by overnight incubation in primary antibody at 1:5000 **(Millipore Cat# ABE457, RRID:AB_2631318)** suspended in blocking buffer. The next day, tissue was washed 3x with 0.1M PBS and then incubated with a solution containing 1:500 Biotin-SP-conjugated Donkey anti-rabbit secondary antibody **(Jackson ImmunoResearch Labs Cat# 711-065-152, RRID:AB_2340593)**. After washing, the antibodies were detected using Streptavidin-conjugated Cy3(1:500), **(Jackson ImmunoResearch Labs Cat# 016-160-084, RRID:AB_2337244)** or Cy5(1:250), **(Jackson ImmunoResearch Labs Cat# 016-170-084, RRID:AB_2337245)**. Finally, sections were counterstained with DAPI (1:10000, Life Technologies) for 15 minutes and mounted on slides (Vectashield mounting medium). Slides were imaged using an Olympus fluorescence virtual slide scanning microscope. For c-fos quantification, 35 μm z-stacks were acquired at 20x magnification. ROIs were chosen in the vCA1 either beneath the optic fiber tip (optogenetic experiments) or that contained AMY projecting neurons (ctb experiment). Fluorescent images were imported into FIJI converted to grayscale and separated by channel. Fluorescent label was marked on each channel independently using the FIJI cell counter tool and the macro metamorph emulator (© 2005 Fabrice P. Cordelières). Overlap (as in ctb experiment) was determined by superimposing the markers from one channel onto another and counting the number of overlapping markers. For any experiments estimating the percent of cells expressing label out of the total number of cells per area, the 3D Objects Counter tool in FIJI was used to estimate the number of DAPI stained nuclei in each area by dividing the obtained volume by the average single nucleus volume for the animal/area.

### 2.5 Virus constructs

The following constructs (AAV2, serotype 5) were packaged by the Vector Core at the University of North Carolina: AAV5 -CaMKII-hChR2(E123T/T159C)-p2A-EYFP-WPRE (Karl Deisseroth) had a titer of 3.6×10^12 - 4.1×10^12 viral particles/ml. AAV5-EF1a-DIO-hChR2 (E123T/T159C) p2A-mCherry (Deisseroth) had a titer of 4.10e^12 virus molecules/ml. AAV5-CAG-FLEX-tdTomato (Boyden) had a titer of 4.8e^12 viral particles/ml. The AAVrg-cre-EBFP plasmid was purchased from Addgene (catalog# 51507) and packaged by the UC Davis Vector Core with a titer of 7.63^12 GC/ml.

### 2.6 Slice preparation for electrophysiological recordings

Mice (postnatal week 6-7; both sexes) were anesthetized through intraperitoneal injection of an anesthetic cocktail (ketamine: 10 mg/kg; xylazine: 1 mg/kg; acepromazine: 0.1 mg/kg) and transcardially perfused with ice-cold artificial CSF (aCSF; in mM: 127 NaCl, 2.5 KCl, 1.25 NaH_2_PO_4_, 25 NaHCO_3_, 1 MgCl_2_, 2 CaCl_2_, 25 glucose; supplemented with 0.4 sodium ascorbate and 2 sodium pyruvate; ~310 mOsm). Brains were rapidly removed, blocked, and placed in choline slurry (110 choline chloride, 25 NaHCO_3_, 25 glucose, 2.5 KCl, 1.25 NaH_2_PO_4_, 7 MgCl_2_, 0.5 CaCl_2_, 11.6 sodium ascorbate, 3.1 sodium pyruvate; ~310 mOsm). Coronal sections (250 μm) containing ventral hippocampus (vCA1) or basal amygdala (BA) were cut on a vibratome (Leica VT1200S) and transferred to an incubation chamber containing aCSF at 32°C for 25 min before moving to room temperature until used for recordings. All solutions were bubbled with 95% O_2_-5% CO_2_ continuously. Chemicals were from Sigma.

### 2.7 Patch-clamp recordings

For recordings, slices were mounted onto glass coveslips coated with poly-l-lysine and placed in a submersion chamber perfused with aCSF (2 ml/min) at 30-32 °C. Loose on-cell patch-clamp recordings were made from visually identified cells in vCA1 or BA using borosilicate glass pipettes (3–5 MΩ) filled with 150 mM NaCl. This configuration does not perturb the intracellular milieu of the recording cell. vCA1 pyramidal neurons were identified based on position and shape and were selected for ChETA-EYFP expression. BA primary neurons (PNs) were identified based on size (> 15 μm) and firing rate (< 20 Hz) (Sosulina et al., 2006; Bazelot, 2015). Recordings were performed in voltage clamp mode by setting the pipette potential to obtain 0 pA of membrane current (Perkins, 2006) and were acquired in pClamp11 using a Multiclamp 700B amplifier (Molecular Devices). Recordings were digitized at 20 kHz with a Digidata 1550 digitizer (Molecular Devices), and low-pass filtered at 8 kHz. Optical stimulation of ChETA-expressing hippocampal pyramidal neurons in vCA1 or their axons in BA was performed under a 60x water immersion lens (1.0 N.A.) of an Olympus BX51W microscope, using an LED system (Prizmatix UHP or Excelitas X-cite; max power of 3 mW at lens tip) mounted on the microscope and driven by a Master9 stimulator (AMPI). Stimulation consisted of 5 10-ms pulses of 488 nm light delivered at various frequencies, as indicated. Each protocol was repeated at least 5 times per stimulation frequency with an inter-trial interval of 30 s [to allow for opsin recovery (Lin, 2011)]. Pulses of increasing power were delivered until an action potential was triggered. Above threshold values (~1.5-2x threshold) were used for experiments. For vCA1 axonal stimulation in BA, higher values were also tested to examine whether more than 1 spikes could be synaptically evoked in BA PNs at 20 Hz. The GABA_A_ receptor blocker bicuculine (20 μM) was washed in during BA recordings, as indicated, for 6 min before resuming stimulation.

### 2.8 Data analysis

For the immediate-early gene and behavioral experiments, group differences were analyzed with ANOVAs and Bonferroni post-hoc tests. All statistics were run in GraphPad Prism (2018) software.

For electrophysiology experiments, data were analyzed with custom-made tools in MATLAB (Mathworks). Spike probability was quantified as the probability of an action potential being evoked during repetitions of the same stimulation regime. For vCA1 recordings, an action potential was considered as evoked if it occurred within a time window of 10 ms from pulse onset (i.e., during the pulse). For BA PN recordings, an action potential was considered as evoked if it occurred within a time window of 15 ms from pulse onset (the longer time window was used to account for synaptic delays). Baseline spike probability was quantified as the average probability of an action potential within 500 randomly selected time windows (10 ms for vCA1; 15 ms for BA) during the 3-s prestimulus baseline. For peri-stimulus time histograms (PSTH), action potentials were counted in 50-ms bins, with time referenced to the start of light pulses.

Permutation tests were used for statistical comparisons of average spike probabilities between conditions (Odén and Wedel 1975). Data were randomly shuffled between conditions 1,000 times, while maintaining the original sample sizes, and the differences between the group averages of observed spike probabilities were compared against the corresponding differences between the group averages of random permutations. The reported p-values indicate the probability that a difference between average spike probabilities equal to or greater than the observed difference could have arisen by chance alone (i.e., due to random sampling).

## 4 Author Contributions

All authors have seen and approved this manuscript. JG and BW designed the research; JG, NV, and RV performed the research; JG, and BW analyzed the data; JG and BW wrote the paper; Fioravante Lab (DF, AD, SJJ, and AP) helped design and performed the electrophysiology experiments. DF and EA analyzed the electrophysiological data, wrote the associated methods, and prepared **Figures 2** and **4**.

---

**Supplementary Figure 1.**
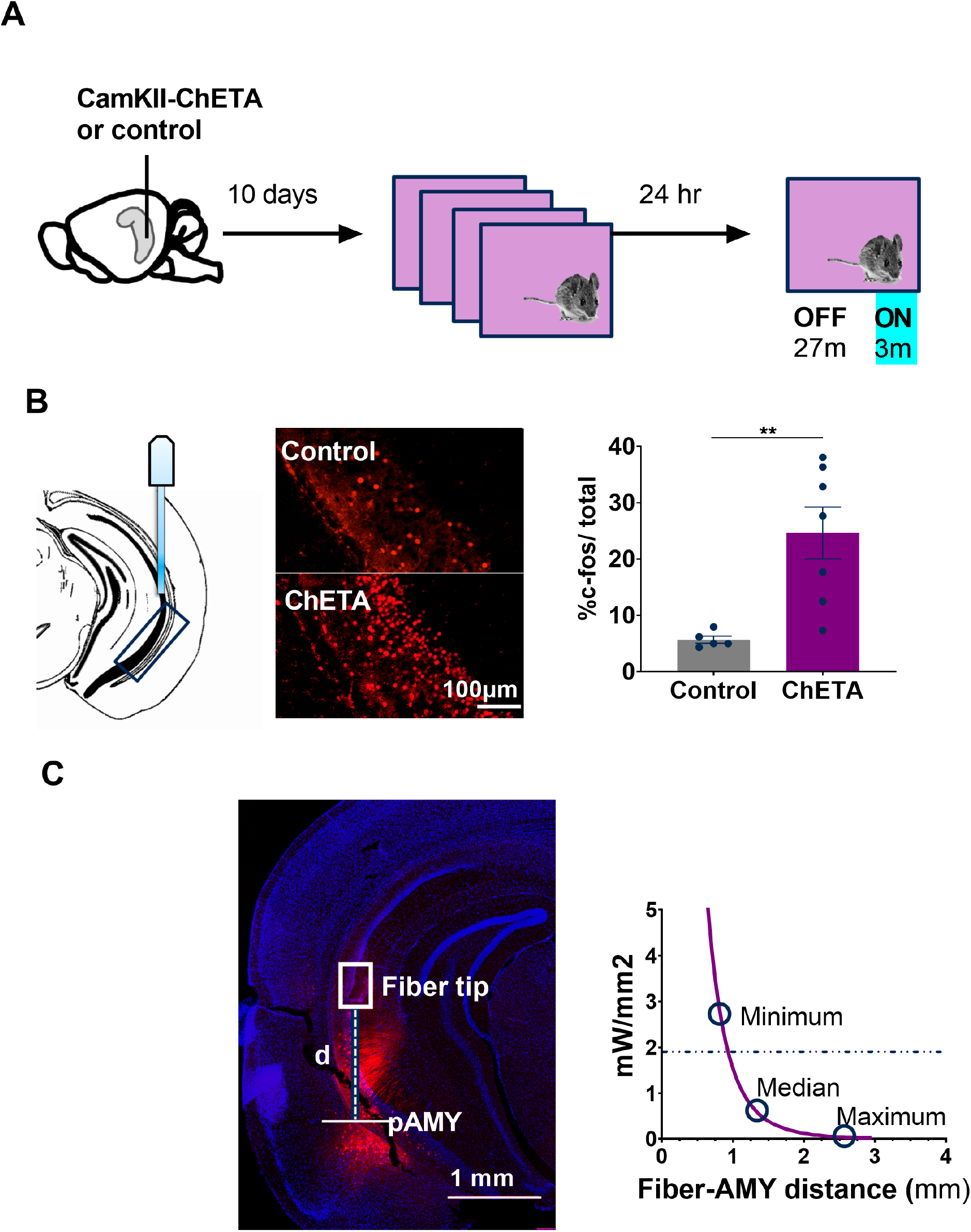

## References

Balogh, S.A., and Wehner, J.A., (2003). Inbred mouse strain differences in the establishment of longterm fear memory. Behav Brain Res 140 (1–2): 97–106. https://doi.org/10.1016/s0166-4328(02)00279-6.

Barot, S.K., Chung, A., Kim, J.J. and Bernstein, I.L. (2009). Functional imaging of stimulus convergence in amygdalar neurons during pavlovian fear conditioning. PLoS ONE 4 (7): 1–8. https://doi.org/10.1371/journal.pone.0006156.

Bazelot, M., Bocchio, M., Kasugai, Y., Fischer, D., Dodson, P.D., Ferraguti, F., and Capogna, M. (2015). Hippocampal theta input to the amygdala shapes feedforward inhibition to gate heterosynaptic plasticity. Neuron 87 (6): 1290–1303. https://doi.org/10.1016/j.neuron.2015.08.024.

Beyeler, A., Chang, C., Silvestre, M., Leveque, C., Namburi, P., Wildes, C.P., and Tye, K.M. (2018). Organization of valence-encoding and projection-defined neurons in the basolateral amygdala. Cell Reports 22 (4): 905–18. https://doi.org/10.1016/j.celrep.2017.12.097.

Cenquizca, L.A. and Swanson, L.W. (2006). Analysis of direct hippocampal cortical field ca1 axonal projections to diencephalon in the rat. The Journal of Comparative Neurology 497 (1): 101–14. https://doi.org/10.1002/cne.20985.

Cenquizca, L.A. and Swanson, L.W. (2007). Spatial organization of direct hippocampal field CA1 axonal projections to the rest of the cerebral cortex. Brain Res Rev 56 (1): 1–26. https://doi.org/10.1016/j.brainresrev.2007.05.002.

Ciocchi, S. Passecker, J., Malagon-Vina, H., Mikus, N. and Klausberger, T. (2015). Selective information routing by ventral hippocampal ca1 projection neurons. Science (New York, N.Y.) 348 (6234): 560–63. https://doi.org/10.1126/science.aaa3245.

Courtin, J., Chaudun, F., R. Rozeske, R.R., Karalis, N., Gonzalez-Campo, C., Wurtz, H., Abdi, A., Baufreton, J., Bienvenu, T.C.M., and Herry, C. (2014). Prefrontal parvalbumin interneurons shape neuronal activity to drive fear expression. Nature 505 (7481): 92–96. https://doi.org/10.1038/nature12755.

Fanselow, M.S., and Dong, H.W. (2010). Are the dorsal and ventral hippocampus functionally distinct structures? Neuron 65. https://doi.org/10.1016/j.conb.2013.11.010.

Fricke, R., and Cowan, W.M. (1978). An autoradiographic study of the commissural and ipsilateral hippocampo-dentate projections in the adult rat. Journal of Comparative Neurology 181 (2): 253–69. https://doi.org/10.1002/cne.901810204.

Hoover, W.B. and Vertes, R.P. (2007). Anatomical analysis of afferent projections to the medial prefrontal cortex in the rat. Brain Structure and Function 212 (2): 149–79. https://doi.org/10.1007/s00429-007-0150-4.

Hübner, C., Bosch, D., Gall, A., Lüthi, A. and Ehrlich, I. (2014). Ex vivo dissection of optogenetically activated mPFC and hippocampal inputs to neurons in the basolateral amygdala: implications for fear and emotional memory. Frontiers in Behavioral Neuroscience 8 (March): 64. https://doi.org/10.3389/fnbeh.2014.00064.

Ishizuka, N., Weber, J. and Amaral, D.G. (1990). Organization of intrahippocampal projections originating from CA3 pyramidal cells in the rat. Journal of Comparative Neurology 295 (4): 580–623. https://doi.org/10.1002/cne.902950407.

Jimenez, J.C., Su, K., Goldberg, A.R., Luna, V.M., Biane, J.S., Ordek, G. et al. (2018). Anxiety cells in a hippocampal-hypothalamic circuit. Neuron 97 (3): 670–83. https://doi.org/10.1016/j.neuron.2018.01.016

Karalis, N., Dejean, C., Chaudun, F., Khoder, S., R Rozeske, Wurtz, H., et al. (2016). 4 Hz oscillations synchronize prefrontal-amygdala circuits during fear behaviour 19 (4): 605–12. https://doi.org/10.1038/nn.4251.4.

Kim, W.B., and Cho, J.H. (2020). Encoding of contextual fear memory in hippocampal–amygdala circuit. Nature Communications 11 (1): 1–22. https://doi.org/10.1038/s41467-020-15121-2.

Kjelstrup, K.B., Solstad, T., Brun, V.H., Hafting, T., Leutgeb, S., Witter, M.P., et al. (2008). Finite scale of spatial representation in the hippocampus. Science 321 (5885): 140–43. https://doi.org/10.1126/science.1157086.

Kjelstrup, K.G., Tuvnes, F.A., Steffenach, H., Murison, R., Moser, E.I., and Moser, M-B. (2002). Reduced fear expression after lesions of the ventral hippocampus. Proceedings of the National Academy of Sciences of the United States of America 99 (16): 10825–30. https://doi.org/10.1073/pnas.152112399.

Komorowski, R.W., Garcia, C.G., Wilson, A., Hattori, S., Howard, M.W., and Eichenbaum, H. (2013). Ventral hippocampal neurons are shaped by experience to represent behaviorally relevant contexts. The Journal of Neuroscience : The Official Journal of the Society for Neuroscience 33 (18): 8079–87. https://doi.org/10.1523/JNEUROSCI.5458-12.2013.

Krueger, J.N., Wilmot, J.H., Teratani-Ota, Y., Puhger, K.R., Nemes, S.E., Crestani, A.P., Lafreniere, M.M., and Wiltgen, B.J. (in press) Amnesia for context fear is caused by widespread disruption of hippocampal activity. Neurobiology of Learning & Memory.

Lesting, J., Narayanan, R.T., Kluge, C., Sangha, S., Seidenbecher, T., and Pape, H.C. (2011). Patterns of coupled theta activity in amygdala-hippocampal-prefrontal cortical circuits during fear extinction. PLoS ONE 6 (6). https://doi.org/10.1371/journal.pone.0021714.

Lovett-Barron, M., Kaifosh, P., Kheirbek, M.A., Danielson, N., Zaremba, J.D., Reardon, T.R. et al. (2014). Dendritic inhibition in the hippocampus suports fear learning. Science 343 (February): 857–64. https://doi.org/10.1126/science.1247485.

Maren, S. and Fanselow, M.S. (1995). Synaptic plasticity in the basolateral amygdala induced by hippocampal formation stimulation in vivo. J Neurosci 15 (11): 7548–64. http://www.ncbi.nlm.nih.gov/pubmed/7472506.

Milanovic, S., Radulovic, J., Laban, O., Stiedl, O., Henn, F., and Spiess, J. (1998). Production of the fos protein after contextual fear conditioning of C57BL/6N mice. Brain Res 784 (1–2): 37–47. http://www.ncbi.nlm.nih.gov/pubmed/9518543.

Moser, E.I., Moser, M-B. and McNaughton, B.L. (2017). Spatial representation in the hippocampal formation: a history. Nature Neuroscience 20 (11): 1448–64. https://doi.org/10.1038/nn.4653.

Moser, M-B. and Moser, E.I. (1998). Functional differentiation in the hippocampus. Hippocampus 8 (6): 608–19. https://doi.org/10.1002/(SICI)1098-1063(1998)8:6<608::AID-HIPO3>3.0.CO;2-7.

Nalloor, R., Bunting, K.M. and Vazdarjanova, A. (2012). Encoding of emotion-paired spatial stimuli in the rodent hippocampus. Frontiers in Behavioral Neuroscience 6 (June): 1–11. https://doi.org/10.3389/fnbeh.2012.00027.

Narayanan, R.T., Seidenbecher, T., Kluge, C., Bergado, J., Stork, O., and Pape, H-C. (2007). Dissociated theta phase synchronization in amygdalo-hippocampal circuits during various stages of fear memory. European Journal of Neuroscience 25 (6): 1823–31. https://doi.org/10.1111/j.1460-9568.2007.05437.x.

Odén, A., & Wedel, H. (1975). Arguments for Fisher’s permutation test. The Annals of Statistics, 3(2), 518–520

O’Keefe, J. and Dostrovsky, J. (1971). The hippocampus as a spatial map. Preliminary evidence from unit activity in the freely-moving rat. Brain Research 34 (1): 171–75. http://www.ncbi.nlm.nih.gov/pubmed/5124915.

O’keefe, J, and Speakman, A. (1987). Single unit activity in the rat hippocampus during a spatial memory task. Exp Brain Res 68: 1–27. https://link.springer.com/content/pdf/10.1007%2FBF00255230.pdf.

Owen, E. H., Logue, S.F., Rasmussen, D.L. and Wehner, J.M. (1997). Assessment of learning by the morris water task and fear conditioning in inbred mouse strains and F1 hybrids: implications of genetic background for single gene mutations and quantitative trait loci Analyses. Neuroscience 80 (4): 1087–99. https://doi.org/10.1016/S0306-4522(97)00165-6.

Padilla-Coreano, N., Bolkan, S.S., Pierce, G.M., Dakota R., Blackman, W.D. Hardin, A. Garcia-Garcia, A.L. et al. (2016). Direct ventral hippocampal-prefrontal input is required for anxiety-related neural activity and behavior. Neuron 89 (4): 857–66. https://doi.org/10.1016/j.neuron.2016.01.011.

Pelletier, J.G., Likhtik, E., Filali, M., and Paré, D. (2005). Lasting increases in basolateral amygdala activity after emotional arousal: implications for facilitated consolidation of emotional memories. Learning & Memory (Cold Spring Harbor, N.Y.) 12 (2): 96–102. https://doi.org/10.1101/lm.88605.2.

Radulovic, J., Kammermeier, J. and Joachim Spiess. (1998). Relationship between fos production and classical fear conditioning : effects of novelty, latent inhibition, and unconditioned stimulus preexposure. 18 (18): 7452–61.

Ramirez, S., Liu, X., Lin, P.A., Suh J., Pignatelli M., Redondo, R.L. et al. (2013). Creating a false memory in the hippocampus. Science (New York, N.Y.) 341 (6144): 387–91. https://doi.org/10.1126/science.1239073.

Ryan, T.J., Roy, D.S., Pignatelli, M., Arons, A., Tonegawa, S. (2015). Engram cells retain memory under retrograde amnesia. Science 348 (6238): 1007–14. https://science.sciencemag.org/content/348/6238/1007.

Seidenbecher, T., Laxmi, R.T., Stork, O., and Pape, H-C. (2003). Amygdalar and hippocampal theta rhythm synchronization. Science 846 (2003): 846–51. https://doi.org/10.1126/science.1085818.

Strange, B.A., Witter, M.P., Lein, E.S., and Moser, E.I. (2014). Functional organization of the hippocampal longitudinal axis. Nature Publishing Group 15 (10): 655–69. https://doi.org/10.1038/nrn3785.

Sutherland, R.J., O’Brien, J. and Lehmann, H. (2008). Absence of systems consolidation of fear memories after dorsal, ventral, or complete hippocampal damage. Hippocampus 18 (7): 710–18. https://doi.org/10.1002/hipo.20431.

Swanson, L.W., Wyss, J.M. and Cowan, W.M. (1978). An autoradiographic study of the organization of intrahippocampal association pathways in the rat. The Journal of Comparative Neurology 181 (4): 681–715. https://doi.org/10.1002/cne.901810402.

Tanaka, K.Z., He, H., Anupratap, T., Niisato, K., Huang, A.J.Y. and McHugh, T.J. (2018). The hippocampal engram maps experience but not place. Science 361 (6400): 392–97. https://doi.org/10.1126/science.aat5397.

Wilmot, J.H, Graham, J.A., LaFreniere, M.M., Puhger, K. and Wiltgen, B.J. (2018) Altered immediate early gene expression in fos-tTA transgenic mice. Society for Neuroscience Poster, 331.26.

Wilmot, J.H., Puhger, K. and Wiltgen, B.J. (2019). Acute disruption of the dorsal hippocampus impairs the encoding and retrieval of trace fear memories. Frontiers in Behavioral Neuroscience 13 (May): 1–9. https://doi.org/10.3389/fnbeh.2019.00116.

Wiltgen, B.J., Sanders, M.J., Anagnostaras, S.G., Sage, J.R. and Fanselow, M.S. (2006). Context fear learning in the absence of the hippocampus. J Neurosci 26 (20): 5484–91. https://doi.org/10.1523/JNEUROSCI.2685-05.2006

Wolff, S.B.E., Gründemann, J., Tovote, P., Krabbe, S., Jacobson, G.A., Müller, C., et al. (2014). Amygdala interneuron subtypes control fear learning through disinhibition. Nature 509 (7501): 453–58. https://doi.org/10.1038/nature13258.

Xu, C., Krabbe, S., Gründemann, J., Botta, P., Fadok, J.P., Osakada, F., et al. (2016). Distinct hippocampal pathways mediate dissociable roles of context in memory retrieval. Cell, 1–12. https://doi.org/10.1016/j.cell.2016.09.051.

